# MAP Kinase and mammalian target of rapamycin are main pathways of gallbladder carcinogenesis: Results from bioinformatic analysis of Next Generation Sequencing data from a hospital-based cohort

**DOI:** 10.1101/2022.05.11.491590

**Authors:** Monika Rajput, SatyaVjiay j Chigurupati, Roli Purwar, Mridula Shukla, Manoj Pandey

## Abstract

**Background:** Gallbladder Cancer (GBC) is one of the most common cancers of the biliary tract and the third commonest gastrointestinal (GI) malignancy worldwide. The disease is characterized by the late presentation and poor outcome despite treatment, and hence, newer therapies and targets need to be identified.

**Methods:** The current study investigated various functionally enriched pathways in GBC pathogenesis involving the genes identified through Next Generation Sequencing (NGS). The Pathway enrichment analysis and Gene Ontology (GO) were carried out after NGS, followed by the construction of the protein-protein interaction (PPI) network to discover associations among the genes.

**Results:** Of the thirty-three patients with GBC who were screened through next-generation sequencing (NGS), 27somatic mutations were identified. These mutations involved a total of 14 genes. The p53 and KRAS were commonly found to be mutated, while mutations in other genes were seen in one case each, the mean number of mutations were 1.2, and maximum mutation in a single case (eight) was seen in one case. The bioinformatics analysis identified MAP kinase, PI3K-AKT, EGF/EGFR, and Focal Adhesion PI3K-AKT-mTOR signaling pathways and cross-talk between these.

**Conclusion:** The results suggest that the complex crosstalk between the mTOR, MAPK, and multiple interacting cell signaling cascades can promote GBC progression, and hence, mTOR - MAPK targeted treatment will be an attractive option.

## INTRODUCTION

Gallbladder cancer (GBC) is a rare malignant neoplasm of the biliary tract and is more prevalent in Asia (1-2). According to GLOBOCAN 2018 data, approximately 1.2% of deaths reported in 2018 were due to GBC (3). It is a relatively rare type of cancer with a poor prognosis with a 5-years survival rate of 10-20%, and a lack of symptoms in its early stages compared to other cancers (4-6). The development of GBC progresses through metaplasia, carcinoma, dysplasia, and invasive malignancy over 5-15 years (7). If GBC is detected earlier and managed effectively, it is completely curable (8). Several internal and external factors are associated with GBC development; of these gallstones (9), various lifestyle-related factors (stress, alcohol, diet, menstrual factors) (10-12), xanthogranulomatous cholecystitis (13), biliary duct infection (14-16), metabolism and lipid peroxidation (17-18), Heavy metals and environmental pollution (19-20) etc., play a crucial role.

Surgery is the primary treatment for early disease, while chemotherapy and radiation are the mainstays in advanced and metastatic GBC (21). The use of other approaches, such as immunotherapy, hormone therapy, and targeted therapy, is mostly experimental, with a slight improvement in progression-free survival with no benefit in overall survival (22). With the advent of advanced methods such as whole-genome sequencing (WGS) using NGS or microarray platforms, the origin of genomic research has expanded, and newer approaches are being identified (23). This has also helped me understand the molecular mechanisms and prediction of treatment response and outcome.

GBC appears to arise due to undiscovered successive spontaneous mutations involving tumor suppressor genes, oncogenes, genes involved with angiogenesis, cell growth and development, and microsatellite instability (24-29). So far in GBC, about 1281 genetic mutations have been discovered (30). Another recent study exploring the transcriptome identified over 900 differentially expressed gene (31). In this study, thirty-three cases of GBC were screened through NGS for mutational studies, and the results of mutation profiling were analyzed using bioinformatics tools to understand the biological pathways involved in gallbladder carcinogenesis and identify a suitable targeted therapy.

### Patients and methods

A prospective study was carried out between January 2017 to December 2021. After approval from the institute ethics committee, and obtaining a written informed consent, naïve patients with a proven histological diagnosis of gallbladder cancer were included in the study.

#### Data collection and processing

Comprehensive history and physical examination of the patients were taken, and details were recorded in the preset proforma. Besides hematology and biochemistry, including the tumor marker CA 19-9, an image-guided biopsy was carried out. CT/MRI/MRCP of the abdomen was carried out to measure the tumor dimensions and stage the disease before initiation of treatment. The tumor tissue was studied for expression of gene mutation by Next-Generation Sequencing. All patients were treated as per standard of care and followed until December 2021.

#### GO and Pathway enrichment analysis

WEB-based Gene Set Analysis Toolkit (Webgestalt) (32), an online bioinformatics tool that helps to investigate significant enriched Genes and functional pathways, Wikicancer Pathway analysis and Gene Ontology (GO) were performed by using Webgestalt tool (http://www.webgestalt.org).

#### Network integration and screening of modules

Protein-protein interaction network (PPI) of genes were constructed by using NetworkAnalyst (http://www.networkanalyst.ca.) (33) and Search Tool for the Retrieval of Interacting Genes (STRING) (http://string-db.org/) (34) based on the confidence scores. Further analysis of involved genes was carried out by constructing a gene-gene interaction network using GeneMania online Tool (35). A signaling network was built among the GBC-specific genes.

#### Disease ontology

Further, Disease Ontology (DO) analysis was performed through Gene Set Enrichment Analysis (GSEA) (36) as a plugin of the Webgestalt tool. Gene-Associated Disease Interaction network of 14 genes was constructed through NetworkAnalyst.

## RESULTS

A total of 33 cases underwent NGS analysis; among them, mutations were identified in 17 of the patients, and a total of 27 mutations (mean 1.19 SD 1.7, range 0-8) were identified in 14 genes which were further analyzed (Table 1). The mutations included TP53 (9 cases), KRAS (4 cases), KDR (4 cases), MAP3K1 (4 cases), BRAF (4 cases), PTEN (2 cases), SMAD4 (1 case), NRAS (1 case), CTNNB1 (1 case), EGRF (1 case), PDGFRA (1 case), FBXW7 (1 case), and POLE (1 case) **(Table 1)**.

**Table 1.**
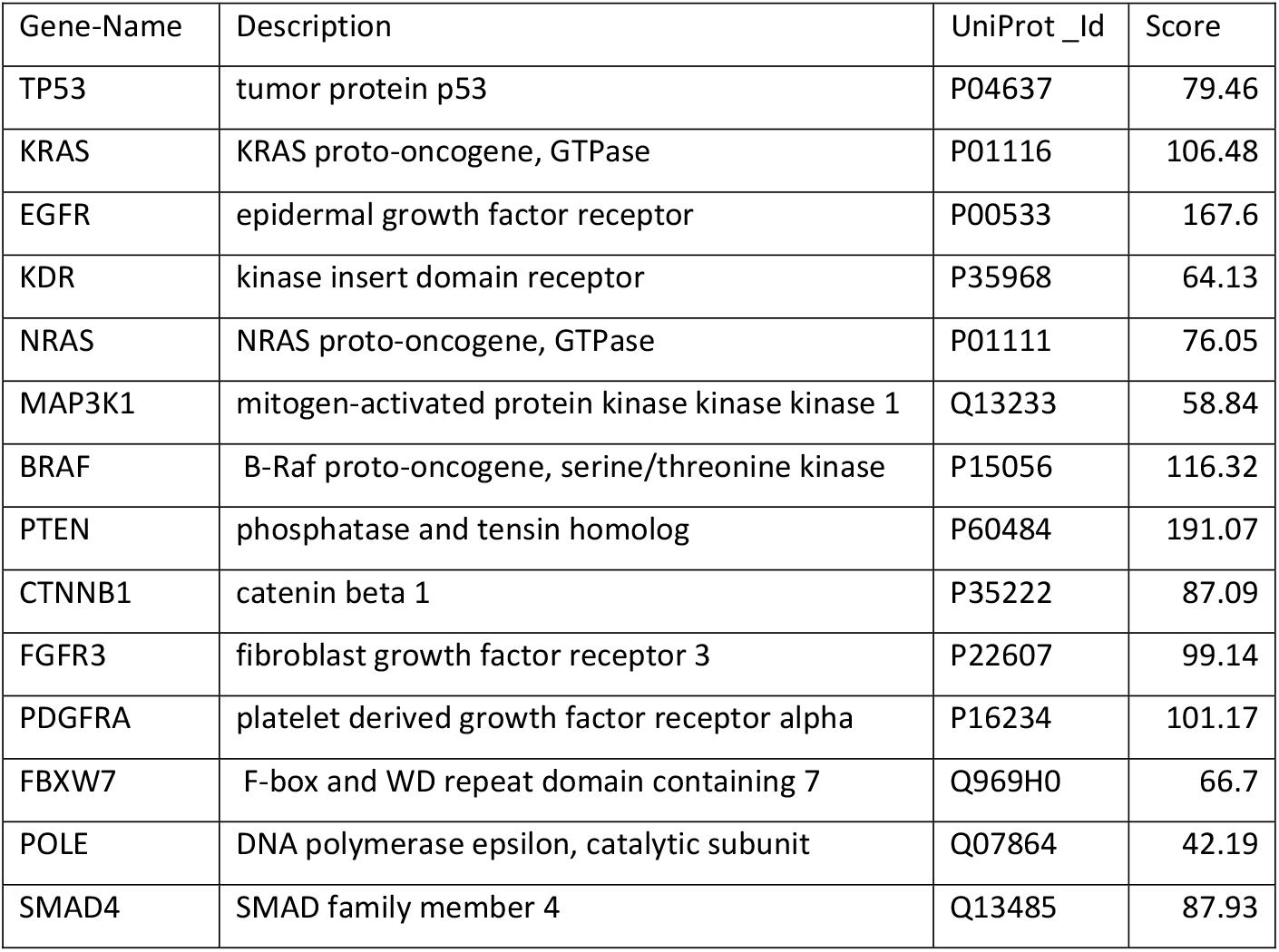
List of 14 significant genes.

| Gene-Name | Description | UniProt_Id | Score |
| --- | --- | --- | --- |
| TP53 | tumor protein p53 | P04637 | 79.46 |
| KRAS | KRAS proto-oncogene, GTPase | P01116 | 106.48 |
| EGFR | epidermal growth factor receptor | P00533 | 167.6 |
| KDR | kinase insert domain receptor | P35968 | 64.13 |
| NRAS | NRAS proto-oncogene, GTPase | P01111 | 76.05 |
| MAP3K1 | mitogen-activated protein kinase kinase kinase 1 | Q13233 | 58.84 |
| BRAF | B-Raf proto-oncogene, serine/threonine kinase | P15056 | 116.32 |
| PTEN | phosphatase and tensin homolog | P60484 | 191.07 |
| CTNNB1 | catenin beta 1 | P35222 | 87.09 |
| FGFR3 | fibroblast growth factor receptor 3 | P22607 | 99.14 |
| PDGFRA | platelet derived growth factor receptor alpha | P16234 | 101.17 |
| FBXW7 | F-box and WD repeat domain containing 7 | Q969H0 | 66.7 |
| POLE | DNA polymerase epsilon, catalytic subunit | Q07864 | 42.19 |
| SMAD4 | SMAD family member 4 | Q13485 | 87.93 |

### Functional enrichment analysis

**The** results of GO enrichment analysis were categorized into three functional categories, i.e. biological processes (BP), molecular function (MF), and cellular components (CC). In the BP, gene enrichment was seen in MAPK cascade, signal transduction by protein phosphorylation, positive regulation of protein phosphorylation, and positive regulation of phosphorylation **(Table 2A)**. In the MF, genes were functionally enriched in transmembrane receptor protein tyrosine kinase activity, transmembrane receptor protein kinase activity, protein-containing complex binding, and MAP kinase activity **(Table 2B)**. In the CC, genes were functionally enriched in the membrane raft, membrane micro domain, membrane region, cell junction, receptor complex, and focal adhesion **(Table 2C)**.

**Table 2(A).**
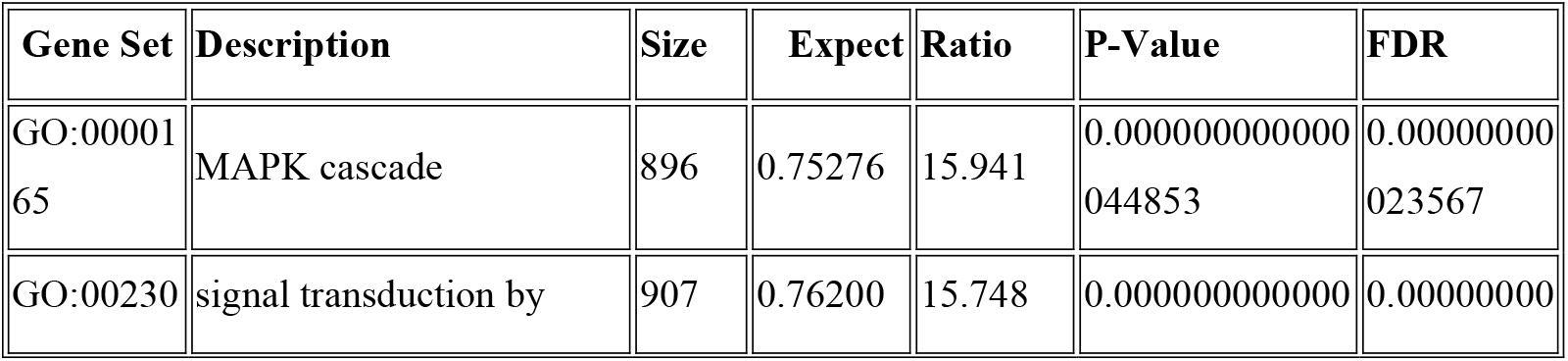
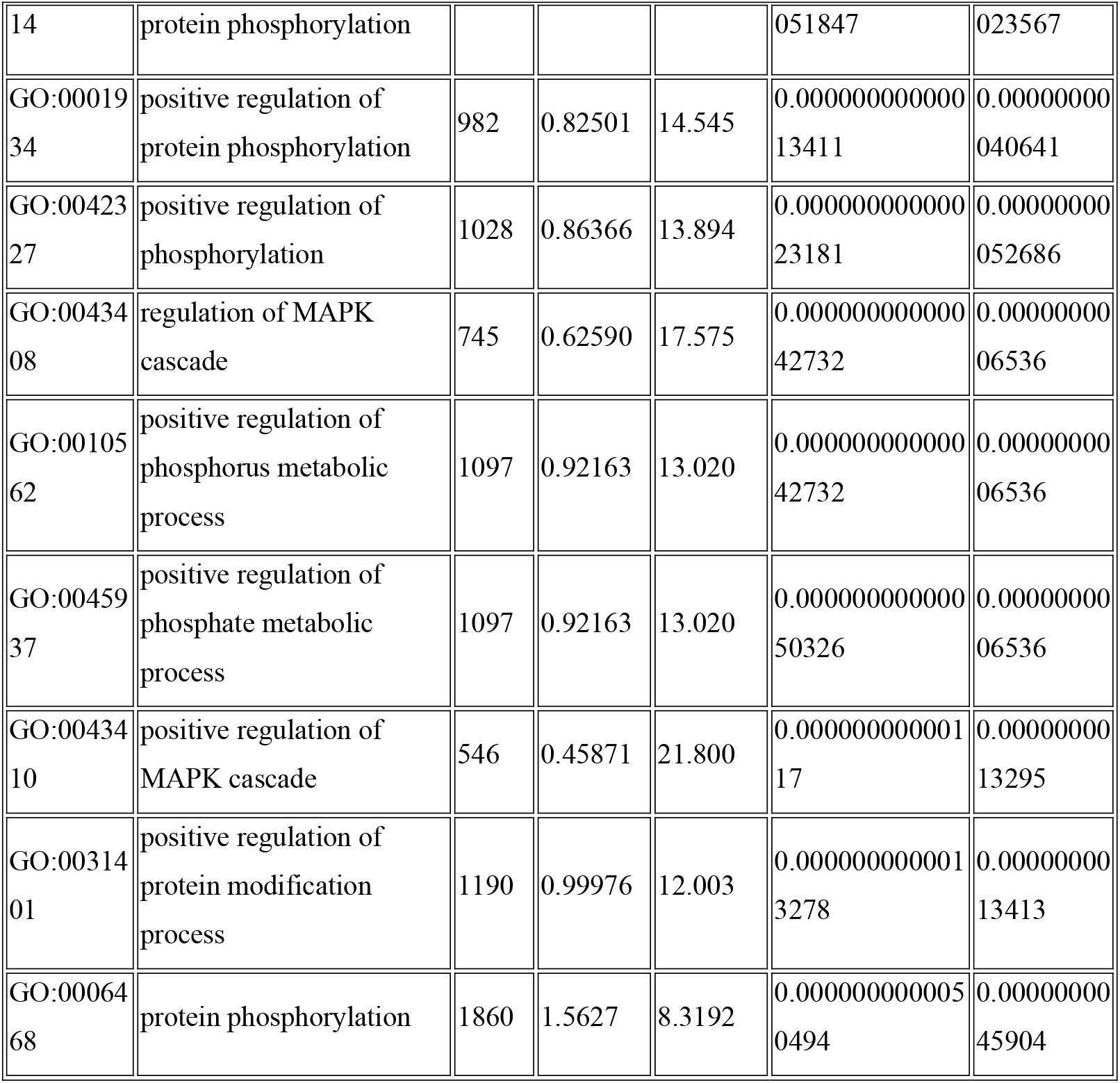
G0: BIOLOGICAL PROCESS. FDR; false discovery rate

**Table 2(B).**
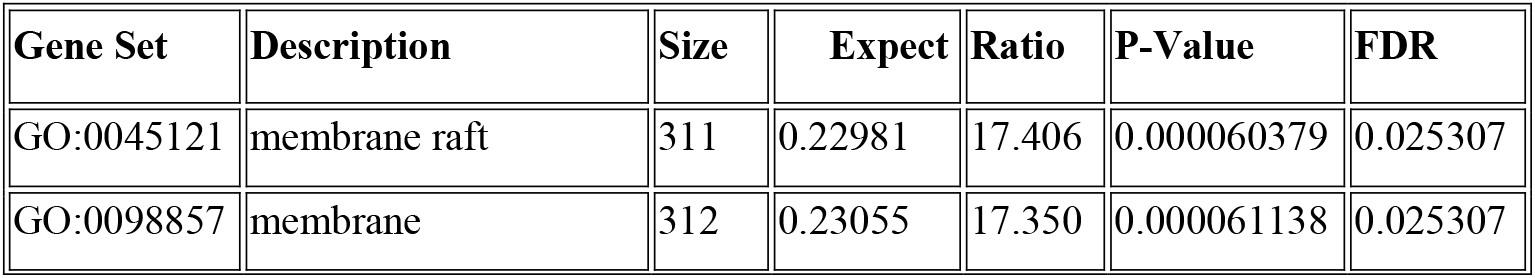
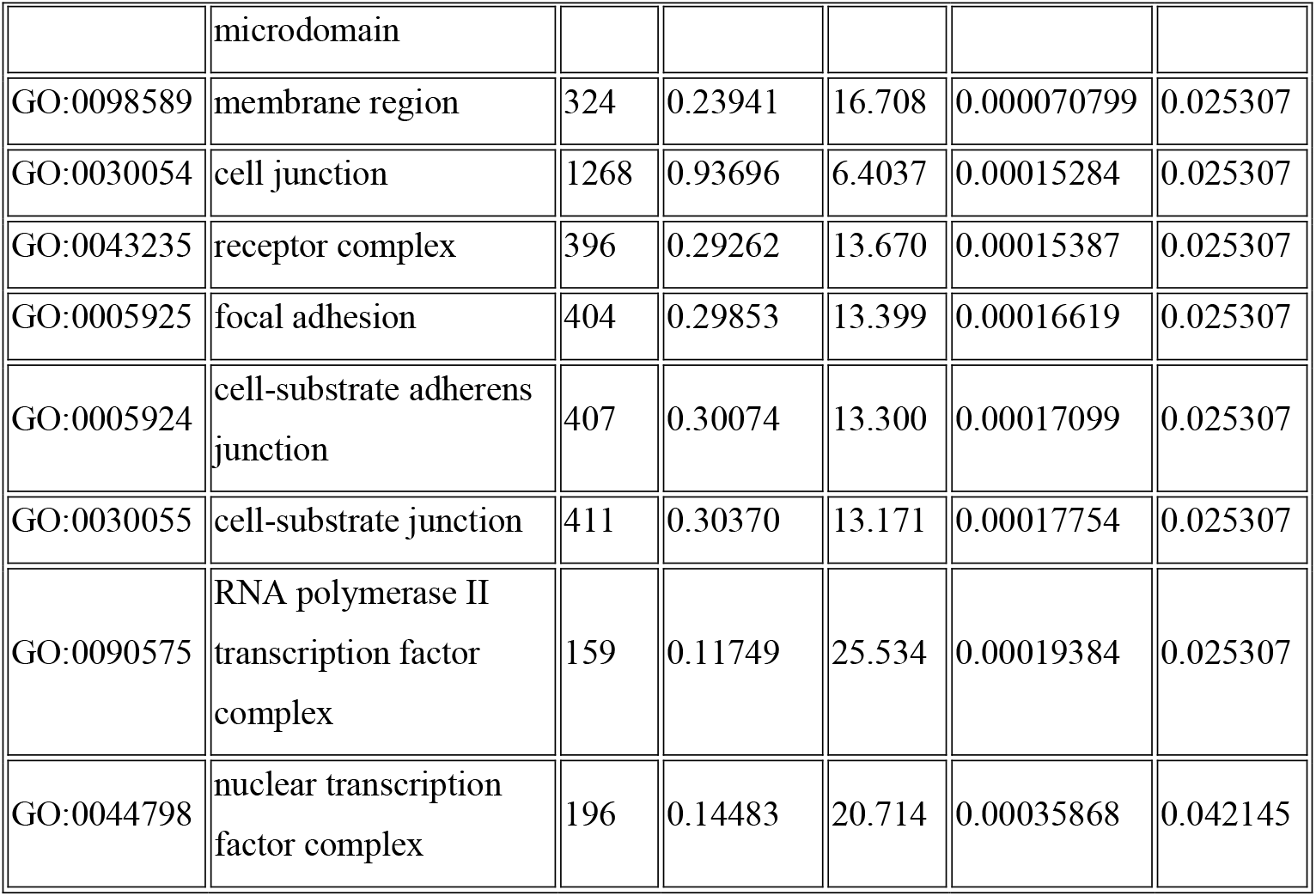
GO: CELLULAR COMPONENT. FDR; false discovery rate

**Table 2(C).** GO: MOLECULAR FUNCTION. FDR; false discovery rate

| Gene Set | Description | Size | Expect | Ratio | P-Value | FDR |
| --- | --- | --- | --- | --- | --- | --- |
| GO:0004714 | transmembrane<br>receptor protein<br>tyrosine kinase activity | 64 | 0.053756 | 74.411 | 0.00000019221 | 0.00035199 |
| GO:0019199 | transmembrane<br>receptor protein kinase<br>activity | 81 | 0.068035 | 58.794 | 0.0000004992 | 0.00035199 |
| GO:0044877 | protein-containing<br>complex binding | 1062 | 0.89201 | 8.9685 | 0.00000056259 | 0.00035199 |
| GO:0004709 | MAP kinase kinase kinase activity | 27 | 0.022678 | 132.29 | 0.0000013635 | 0.00053512 |
| GO:0042802 | identical protein binding | 1696 | 1.4245 | 6.3179 | 0.0000014255 | 0.00053512 |
| GO:0035639 | purine ribonucleoside triphosphate binding | 1786 | 1.5001 | 5.9995 | 0.0000022127 | 0.00066600 |
| GO:0032555 | purine ribonucleotide binding | 1850 | 1.5539 | 5.7920 | 0.0000029824 | 0.00066600 |
| GO:0017076 | purine nucleotide binding | 1865 | 1.5665 | 5.7454 | 0.0000031934 | 0.00066600 |
| GO:0032553 | ribonucleotide binding | 1865 | 1.5665 | 5.7454 | 0.0000031934 | 0.00066600 |
| GO:0019838 | growth factor binding | 138 | 0.11591 | 34.509 | 0.0000042221 | 0.00079249 |

### Pathway enrichment analysis

**Figure 1** shows the ten positive and four negatively related categories according to the false discovery rate (FDR > 0.05). The genes significantly enriched in MAPK, PI3K-AKT, EGF/EGFR, Focal Adhesion, and PI3K-AKT-mTOR signaling pathway. The resulted outcome stipulated that these significant genes were functionally enriched in cancer-associated biological pathways **(Figure 2)**.

**Figure 1.**
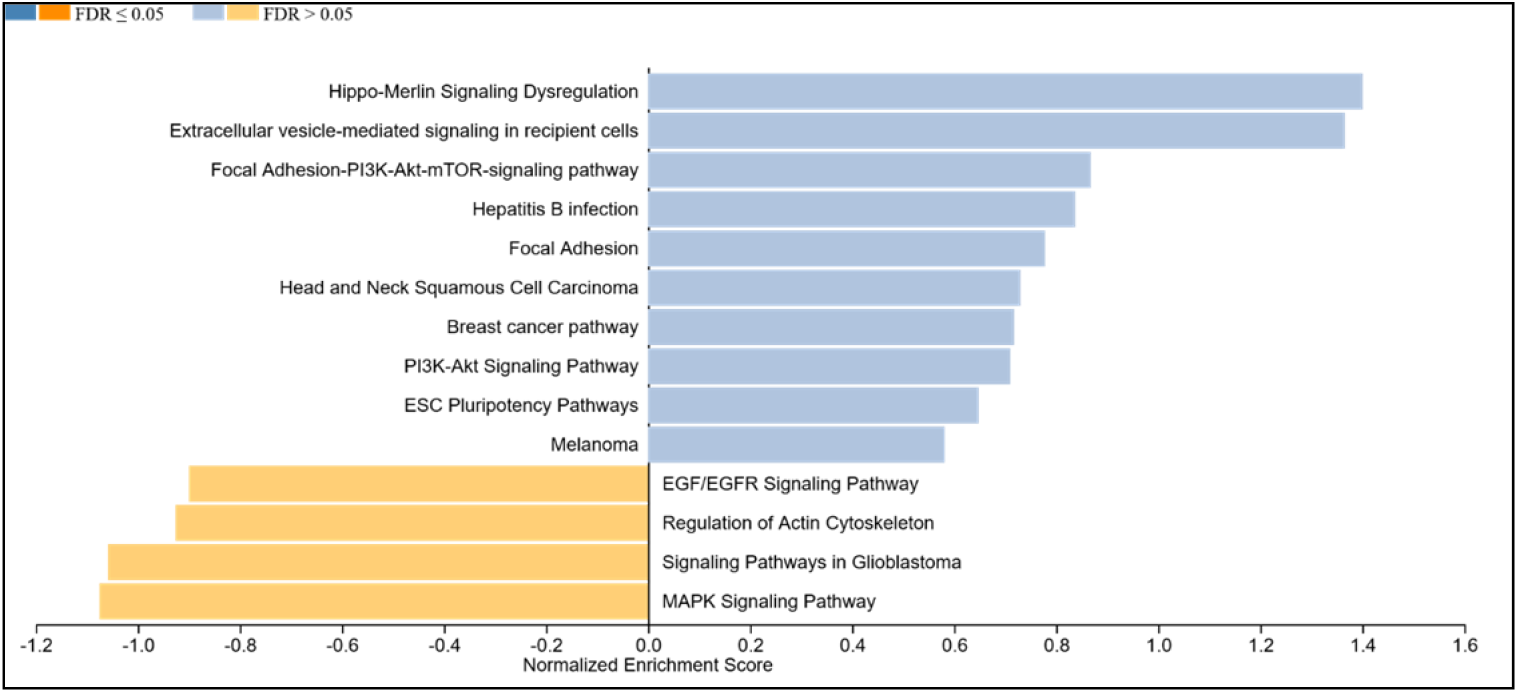
PATHWAY ANALYSIS: WIKICANCER PATHWAY. **10 positive related** categories and **4 negative related** categories are identified as enriched categories.

**Figure 2.**
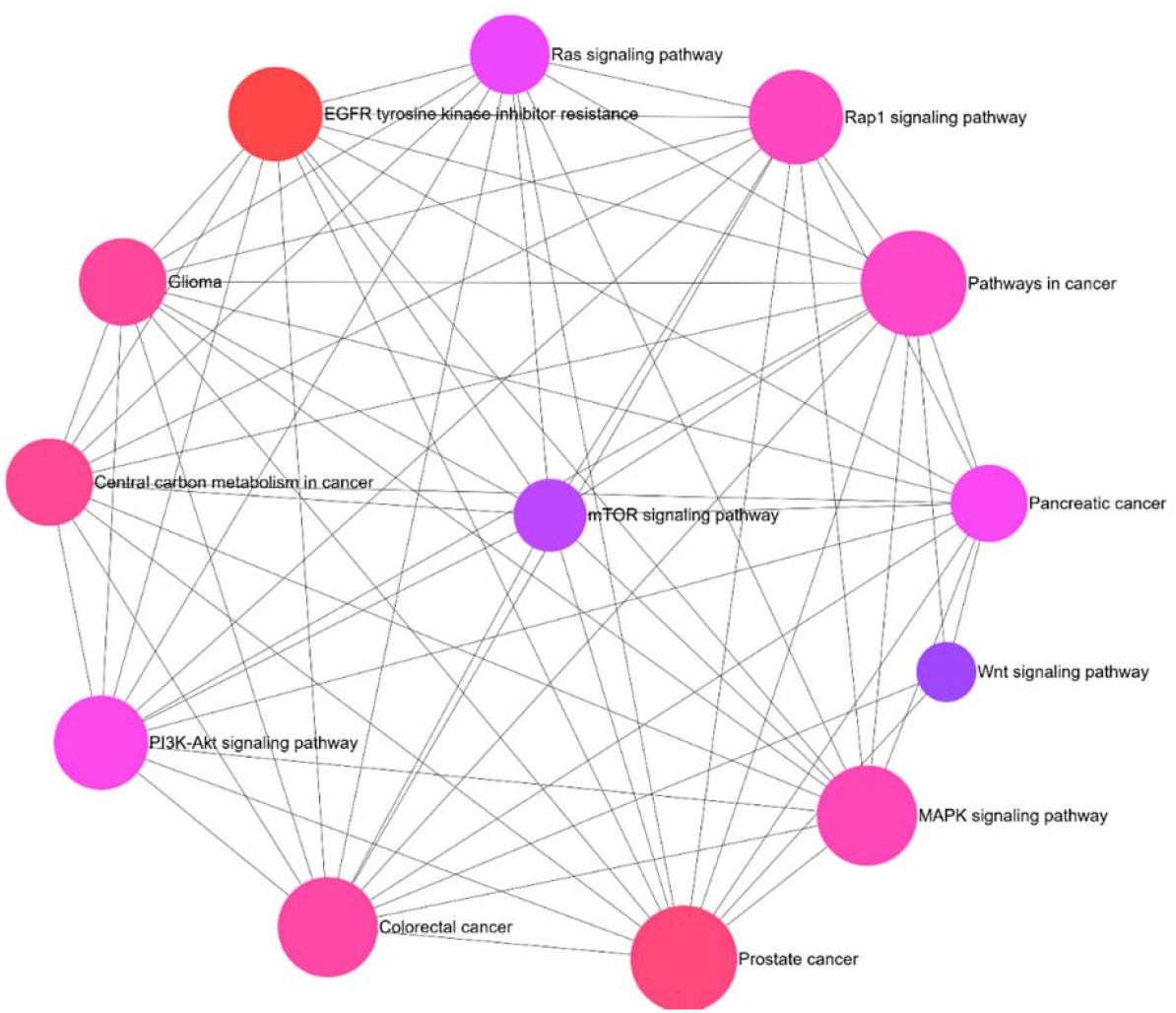
Pathway Network.

### GBC-specific genes functional characterization

The Protein-Protein Interaction (PPI) network of 14 significant genes with 14 nodes and 71 edges was constructed using STRING **(Figure 3)**. Further, several hub genes exhibiting co-expression, predicted, and physical and genetic interaction with multiple genes were identified, and a network was constructed through the GeneMania tool **(Figure 4 and Additional Table 1)**. The NetworkAnalyst Tool built a signaling network through a plugin Signor (https://signor.uniroma2.it) (37) of 13 significant genes, with 503 nodes and 619 edges shown in **(Figure 5)**.

**Figure 3.**
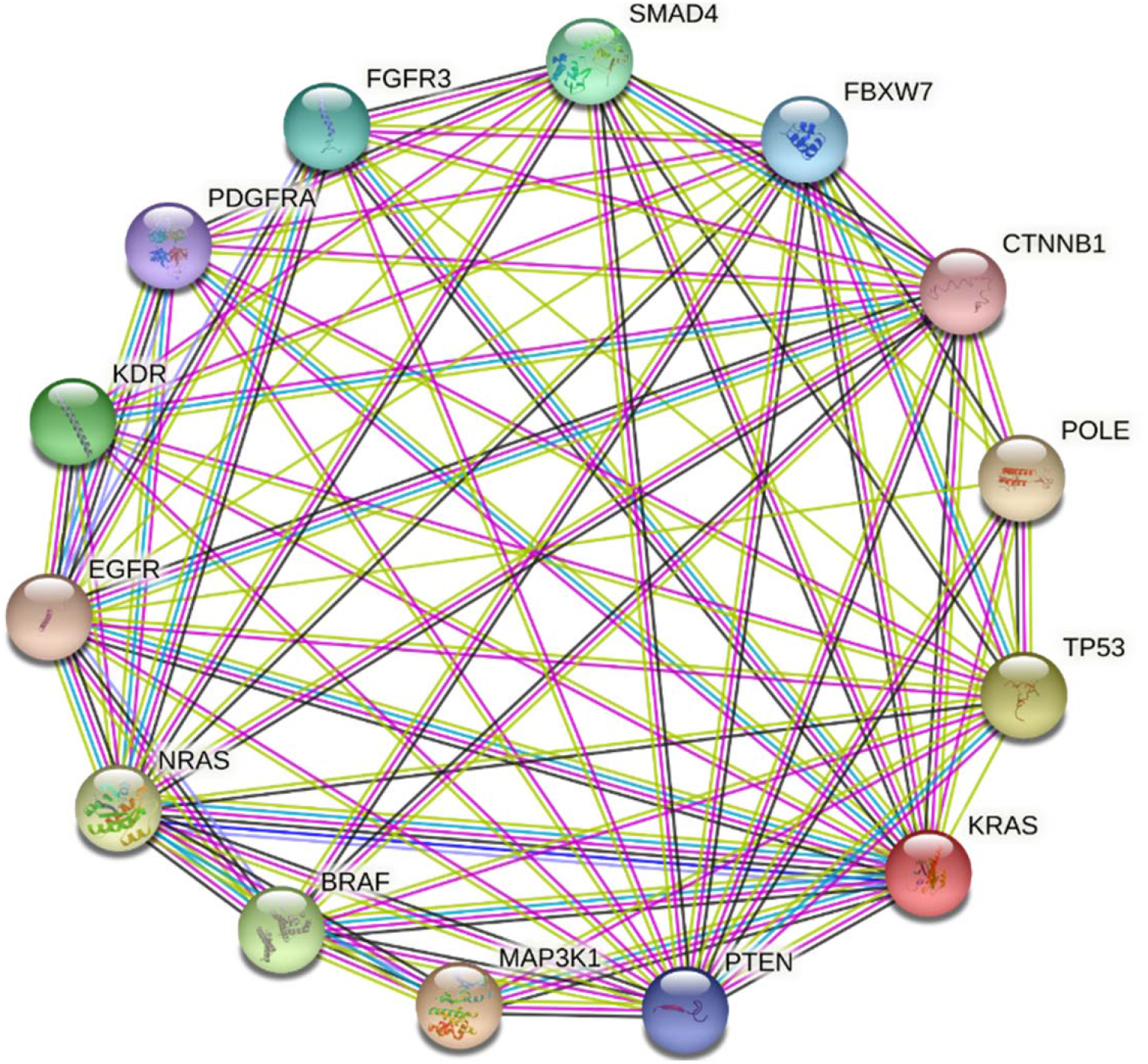
Protein-Protein Interaction network of 14 genes was constructed in STRING. Known Interactions - curated databases – light blue; experimentally determined – pink Predicted Interactions - gene neighborhood – green; gene fusions – red; gene co-occurrence – dark blue Others - Text-mining – yellow; co-expression – black; protein homology – grey

**Figure 4.**
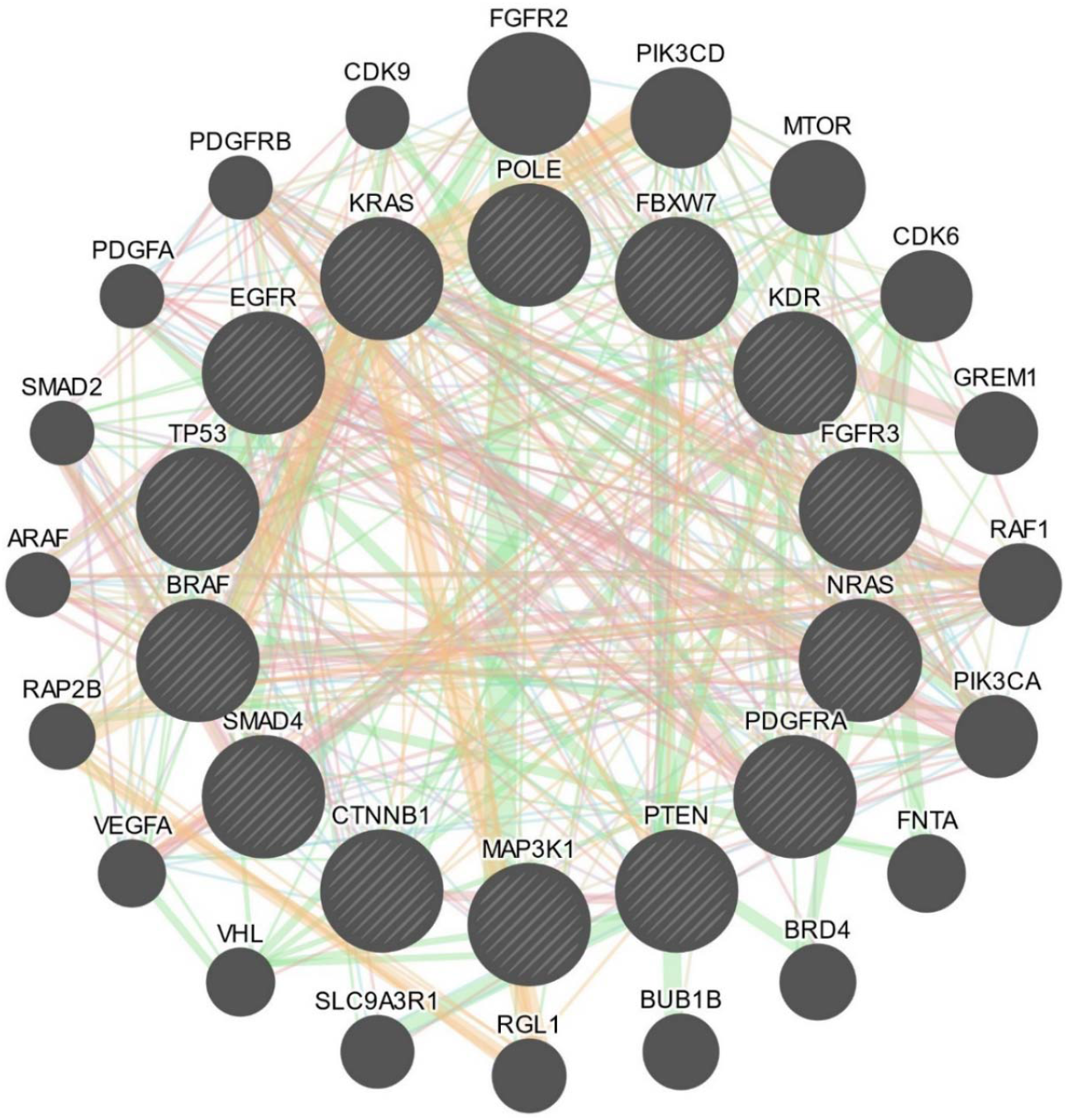
Gene-Gene Interaction network of 14 genes was constructed in GeneMania. Colored edges represented the interaction between the genes; physical interaction- pink; genetic interaction- green; predicted- orange; co-expression- purple; shared protein domain- grey; pathway- light blue.

**Figure 5.**
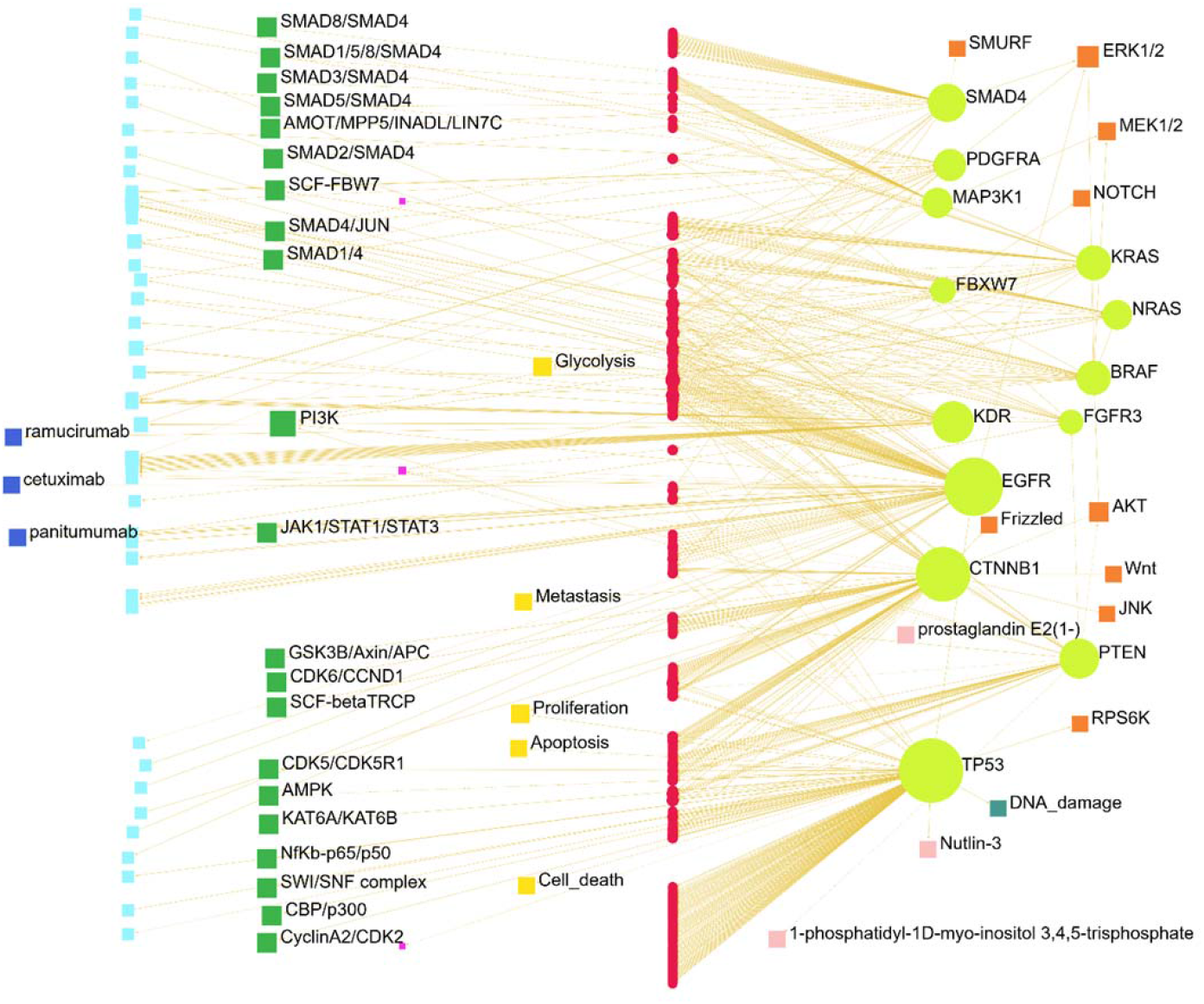
Signaling network. Genes (circle light green), complex (Circle light green), pink (circle dark), chemical (square light blue), protein family (square orange), small molecule (square light pink), stimulus (square blue-green), and phenotype (square yellow).

### Disease ontology (DO)

Further, DO with FDR (> 0.05) was functionally enriched. The enriched DO showed identified genes to be associated with Melanoma, Gastrointestinal Diseases, Gastrointestinal Neoplasms, Carcinoma and Squamous cells, Nervous System Neoplasms, and Breast Neoplasms **(Figure 6)**. The results of the Gene-Disease associated network of significantly enriched six genes in NetworkAnalyst are presented in **(Additional Figure 1)**.

**Figure 6.**
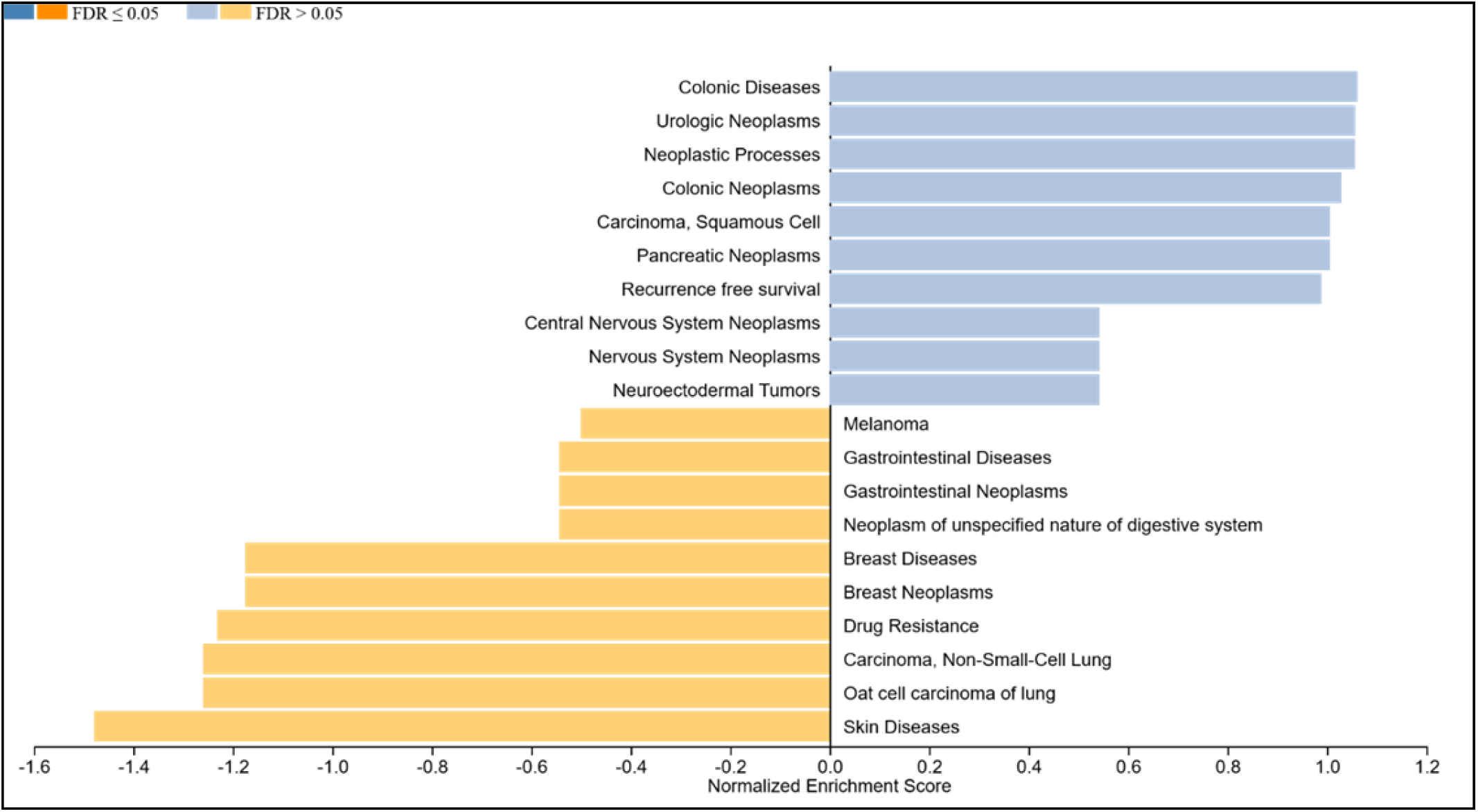
Disease Ontology of 14 genes was constructed through Webgestalt. **10 positive related** categories and **7 negative related** categories are identified as enriched categories.

### Cross talk between mTOR/MAPK signaling pathway

High frequency of the mammalian target of Rapamycin (mTOR) & the mitogen-activated protein kinase (MAPK) signaling pathways variation was observed, including PTEN, AKT, TP53, SMAD4, EGFR, and CTNNB1 **(Figure 7)** a pathway of cross-talk between various identified pathways was constructed by data and text mining.

**Figure 7.**
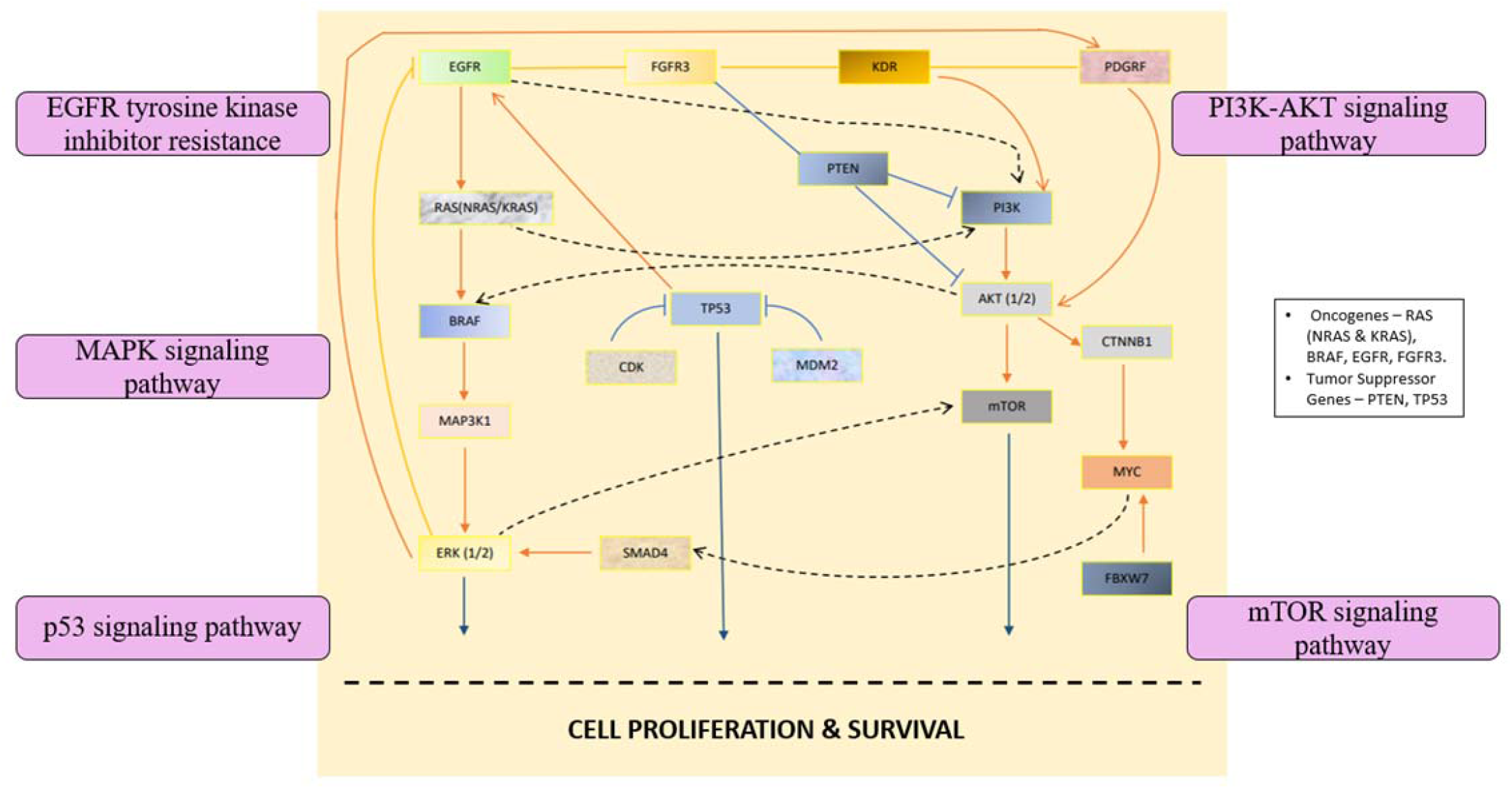
Crosstalk among associated pathways.

## DISCUSSION

Multi-omics characterization of the NGS data from GBC patients identified 14 significant genes and their functional and biological pathways, with MAP kinase and mTOR being the main. Despite recent breakthroughs in surgical procedures and drug development, gallbladder cancer has a dismal long-term prognosis, with a 5-year survival rate ranging from 5% to 13% (38) (39).

Gain-of-function mutations in FGFRs have been described in numerous malignancies, and they play a crucial role in angiogenesis and proliferation (40). To our knowledge, no FGFR3 mutation or amplification has been documented in gallbladder cancer. Hence, the discovery of Fibroblast Growth Factor Receptor (FGFR3) was a novel result in our investigation.

Moreover, the investigation uncovered that the TP53 family is associated with different mutation in TP53, in most of our cases suggestingthat it acts as a mutagenic driver in GBC. TP53 is the commonest gene studied in the gallbladder and extrahepatic biliary tract cancer. TP53 mutations with or without RAS mutations are reported in up to 50% of gallbladder cancer patients (41). No difference is observed in patients with the anomalous junction of the pancreatic, and biliary duct (42). In some areas, it’s higher, while others display lower p53 mutation rates (43). Bolivia reported 50% mutation rates in their patients, all but one patient had a single mutation, while one had three mutations in the same gene. Most of these mutations were on exons 5 and 8 of the gene (44). Eighty single nucleotide variants and 8 indels in 39 genes were identified in their patients with biliary tract cancer, including gallbladder, p53, and KRAS, were the commonest mutations identified in these patients (44). KRAS is a well-known oncogene, commonly mutated in various malignancies (45). Patients with GBC were found to have a mutation in the KRAS gene.

A number of other targets like EGFR, VEGF, BRAF, MAPK, etc., were identified, some of which for the first time. Further, significantly higher identification of mutations in p53 and RAS oncogene signifies that treatment by EGFR antibodies may not be successful in these cases. However, the relatively widespread frequency of MAPK and mTOR signaling pathway mutations (NRAS, BRAF, TP53, AKT, MAPK31, and PTEN) was a remarkable result, opening up possible alternatives for targeted therapy directed against the mTOR pathway. Previous studies have also shown the importance of MAP kinase and mTOR pathways in gallbladder cancer. A study in the gallbladder cancer cell line from typhoid carriers and an animal model from the same cell line showed mTOR as the main pathway of carcinogenesis, leading authors to suggest targeting of mTOR receptors (46). Further experimental studies demonstrated regression of gallbladder cancer by treatment with mTOR inhibitors (47). This was independent of the typhoid carrier state and was demonstrated to be mediated through PIK3CA/AKT/mTOR pathway (48-52). Although the single-phase I study of docetaxel and temsirolimus was limited by severe myelosuppressive toxicity and failed to meet the objectives (53).

Results of the present study, and bioinformatics show cross-talk between various pathways with mTOR, including the EGFR pathway, p53 pathway, and PIK3CA/AKT pathway, suggesting the need to conduct clinical trials on mTOR inhibitors. The results of this study gives a unique insight into gallbladder carcinogenesis, identifies driver oncogenes, and suggest new therapeutic strategies that need to be tested.

## CONCLUSION

The study reports the results of DNA sequencing and demonstrated 14 key genes in gallbladder carcinogenesis, including P53, RAS, EGFR, MAP3K1, PTEN, etc. The analysis also demonstrates that the mTOR and MAPK signaling networks were major pathways in gallbladder carcinogenesis. We suggest that the complex crosstalk between the mTOR, MAPK, and multiple interacting cell signaling cascades promotes gallbladder carcinogenesis by activating cell division. This suggests that mTOR inhibitors are an attractive option in the treatment of treatment gallbladder cancer, and this needs to be tested in clinical trials.

## Authors Contribution

MR: Conduct of the study, bioinformatics analysis and interpretation and preparation of the draft manuscript

VJC: Collection of the data, design of study, interpretation of results and preparation of manuscript

RP: data collection, interpretation of results and preparation of manuscript

MS: Interpretation of pathological and molecular results, preparation of manuscript

MP: concept and design, interpretation of results, Editing of the final manuscript

All authors read and approved final manuscript for publication

## Ethical approval and Consent

The study was approved by the Institute Ethics committee vide approval letter no Dean/EC/2020/2045 dated 18.7.2020 written informed consent was taken from all patients participating the study

## Consent to Publish

Not applicable

## Acknowledgement

None

## Abbreviations

GO: Gene Ontology
PPI: Protein-protein interaction network
Webgestalt: WEB-based Gene Set Analysis Toolkit
STRING: Search Tool for the Retrieval of Interacting Genes
mTOR: the mammalian target of Rapamycin
MAPK: the mitogen-activated protein kinase signaling pathways

## Legends for figures

**Additional Figure 1.**
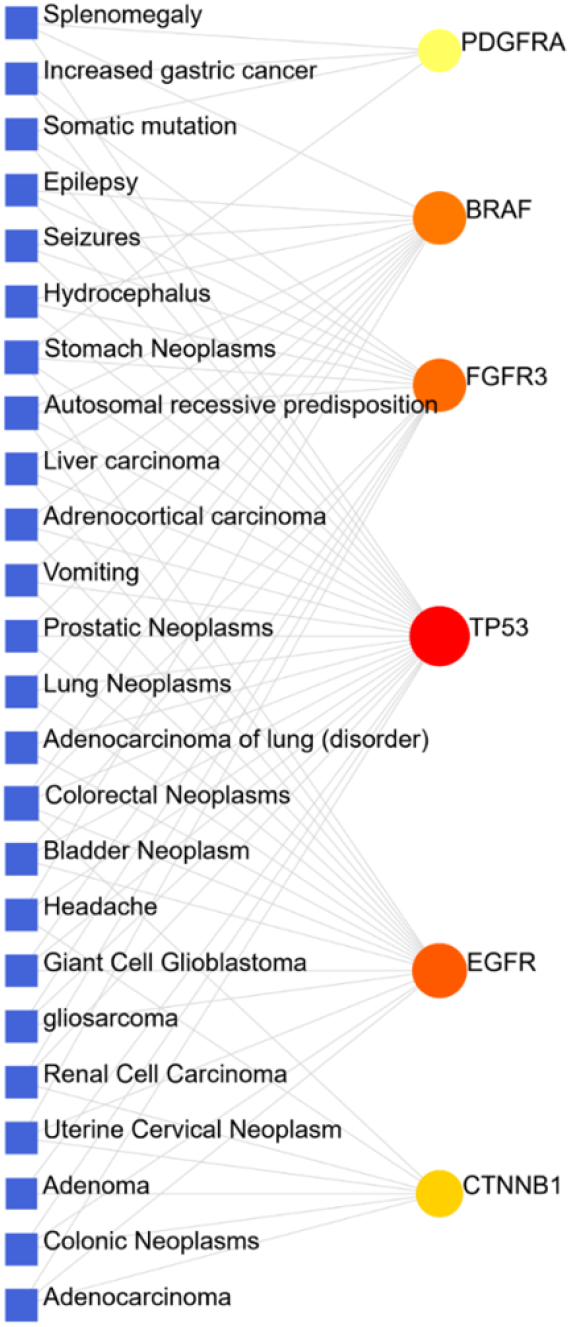
Gene-Associated disease Interaction network of 14 genes was constructed through NetworkAnalyst. Gene-Disease associated network of significantly enriched 6 genes was constructed in NetworkAnalyst. Here, colored circular dots represent genes whereas a blue-colored square box represents associated diseases.

**Additional Table 1.**
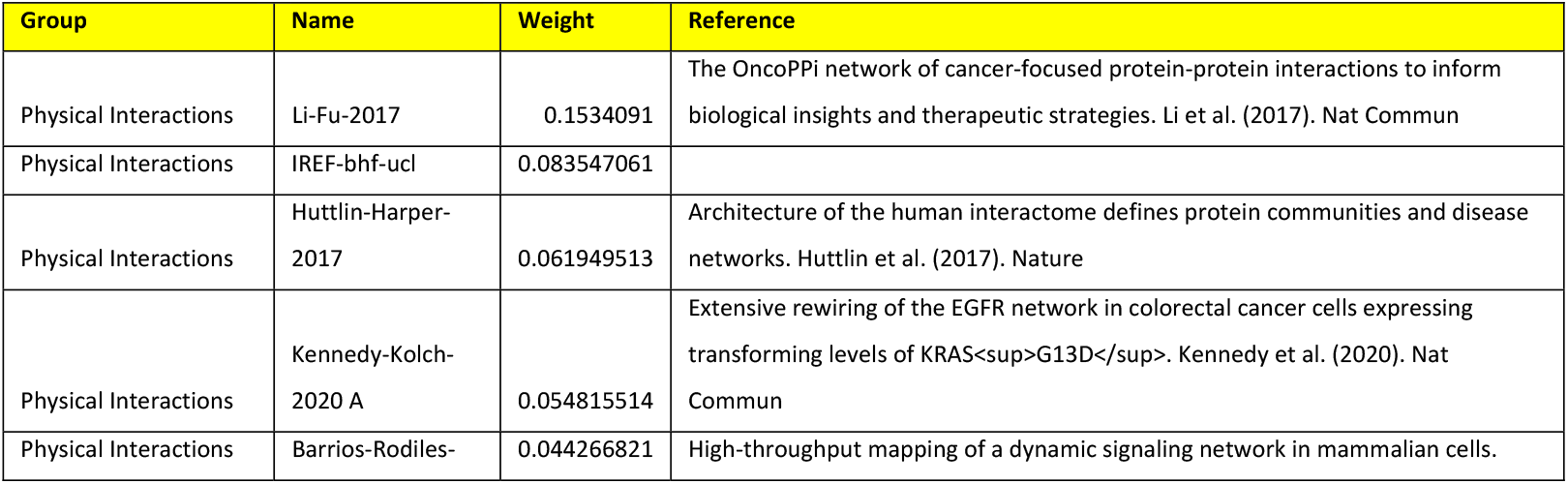
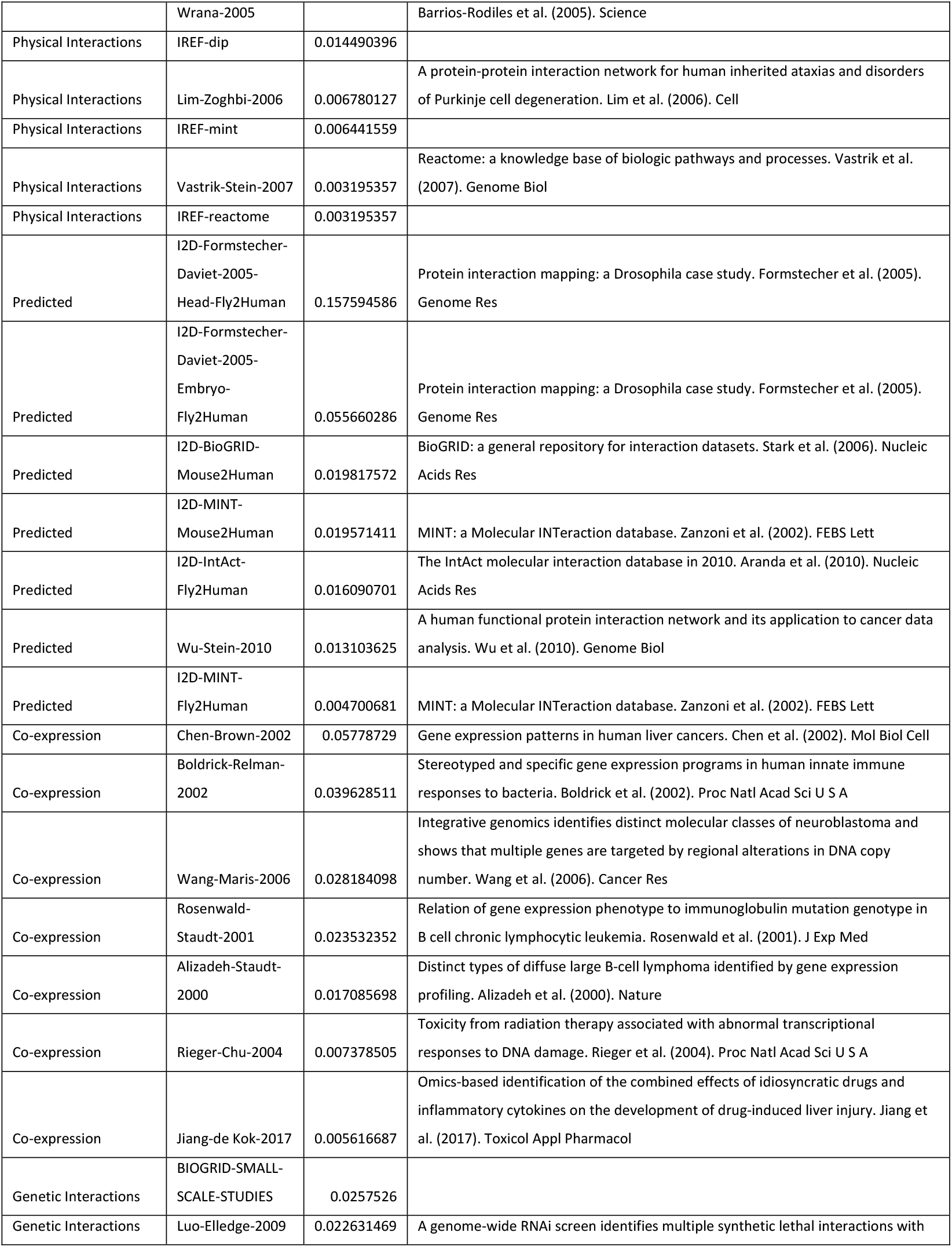
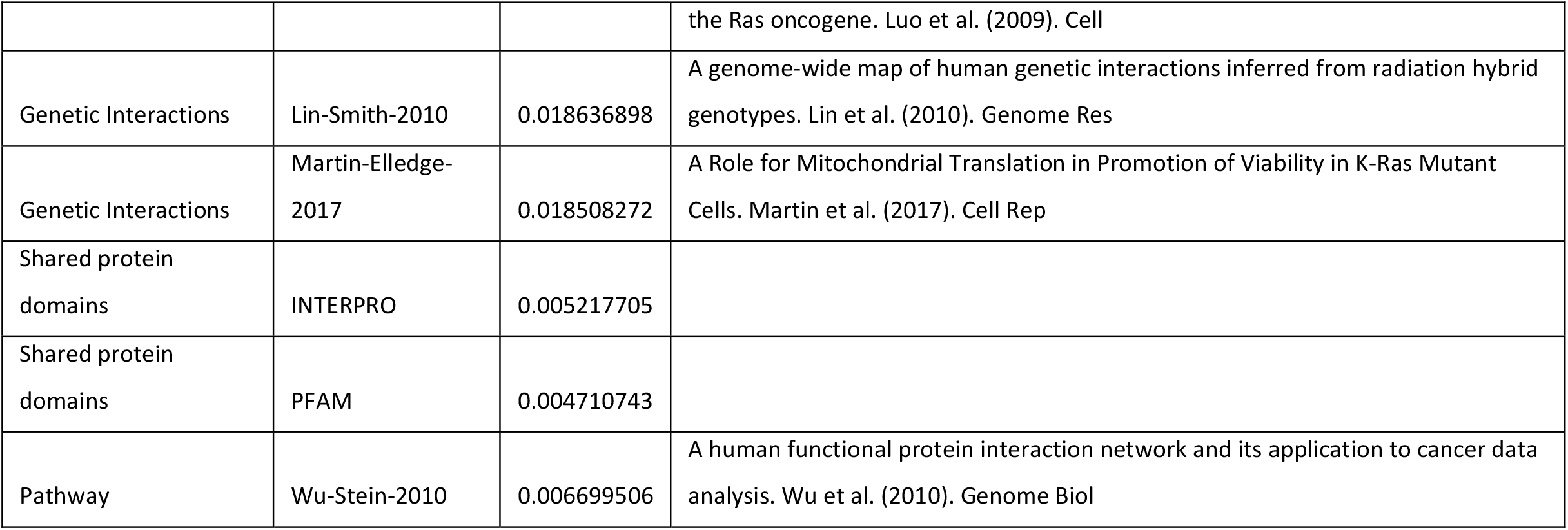
Genemania Network.

## REFERENCES

1. Mehrotra R, Tulsyan S, Hussain S, Mittal B, Singh Saluja S, Singh S, Tanwar P, Khan A, Javle M, Hassan MM, Pant S, De Aretxabala X, Sirohi B, Rajaraman P, Kaur T, Rath GK. Genetic landscape of gallbladder cancer: Global overview. Mutat Res Rev Mutat Res. 2018 Oct-Dec;778:61–71. doi: 10.1016/j.mrrev.2018.08.003. Epub 2018 Aug 23. PMID: 30454684.

2. Jin H, Cui M. Gene silencing of heparanase results in suppression of invasion and migration of gallbladder carcinoma cells. Biosci Biotechnol Biochem. 2018 Jul;82(7):1116–1122. doi: 10.1080/09168451.2018.1456316. Epub 2018 Mar 29. PMID: 29598788.

3. Rawla P, Sunkara T, Thandra KC, Barsouk A. Epidemiology of gallbladder cancer. Clin Exp Hepatol. 2019 May;5(2):93–102. doi: 10.5114/ceh.2019.85166. Epub 2019 May 23. PMID: 31501784; PMCID: PMC6728871.

4. Nemunaitis JM, Brown-Glabeman U, Soares H, Belmonte J, Liem B, Nir I, Phuoc V, Gullapalli RR. Gallbladder cancer: review of a rare orphan gastrointestinal cancer with a focus on populations of New Mexico. BMC Cancer. 2018 Jun 18;18(1):665. doi: 10.1186/s12885-018-4575-3. PMID: 29914418; PMCID: PMC6006713.

5. Akhtar J, Priya R, Jain V, Sakhuja P, Agarwal AK, Goyal S, Polisetty RV, Sirdeshmukh R, Kar S, Gautam P. Immunoproteomics approach revealed elevated autoantibody levels against ANXA1 in early stage gallbladder carcinoma. BMC Cancer. 2020 Dec 1;20(1):1175. doi: 10.1186/s12885-020-07676-6. PMID: 33261560; PMCID: PMC7709428.

6. Horsley-Silva JL, Rodriguez EA, Franco DL, Lindor KD. An update on cancer risk and surveillance in primary sclerosing cholangitis. Liver Int. 2017 Aug;37(8):1103–1109. doi: 10.1111/liv.13354. Epub 2017 Jan 28. PMID: 28028930.

7. Espinoza JA, Bizama C, García P, Ferreccio C, Javle M, Miquel JF, Koshiol J, Roa JC. The inflammatory inception of gallbladder cancer. Biochim Biophys Acta. 2016 Apr;1865(2):245–54. doi: 10.1016/j.bbcan.2016.03.004. Epub 2016 Mar 12. PMID: 26980625; PMCID: PMC6287912.

8. Kanthan R, Senger JL, Ahmed S, Kanthan SC. Gallbladder Cancer in the 21st Century. J Oncol. 2015;2015:967472. doi: 10.1155/2015/967472. Epub 2015 Sep 1. PMID: 26421012; PMCID: PMC4569807.

9. Espinoza JA, Bizama C, García P, Ferreccio C, Javle M, Miquel JF, Koshiol J, Roa JC. The inflammatory inception of gallbladder cancer. Biochim Biophys Acta. 2016 Apr;1865(2):245–54. doi: 10.1016/j.bbcan.2016.03.004. Epub 2016 Mar 12. PMID: 26980625; PMCID: PMC6287912.

10. Pandey M, Singh S, Shukla VK. Diet and gallbladder cancer: A casecontrol study. Eur J Cancer Prevention 2002; 11: 365–8. PMID: 12195163

11. Pandey M, Singh S, Shukla VK. Life-style, parity, Menstrual and reproductive factors and gallbladder cancer. Eur J Cancer Prev 2003; 12: 269–72. PMID: 12883378

12. Pandey M. Risk factors for gallbladder cancer a reappraisal. Eur J Cancer Prev 2003; 12: 15–24. PMID: 12548106

13. Dixit VK, Prakash A, Gupta A, Pandey M, Kumar M, Gautam A, Shukla VK. Xanthogranulomatous cholecystitis. Dig Dis Science 1998; 43(5): 940–2. PMID: 9590403

14. Shukla VK, Singh H, Pandey M, Upadhyaya SK, Nath G. Carcinoma of the gallbladder is it a sequel of typhoid? Dig Dis Sci 2000; 45: 900–3. PMID: 10795752

15. Pandey M, Shukla M. Helicobacter species are associated with possible increase in risk of hepatobiliary cancers. Surgical Oncology 2009; 18:51–56 PMID: 18715780

16. Pandey M, Mishra RR, Dixit R, Jaiswal R, Shukla M, Nath G. Helicobacter bilis in human gallbladder cancer: Results of a case control study and meta analysis. Asia Pacific journal of Epidemiology and prevention 2010; 11: 343–47.PMID: 20843113

17. Pandey M, Shukla VK. Fatty acids, biliary bile acids, lipid peroxidation products and gallbladder cancer: A hypothesis. European J Cancer Prevention 2000; 9:165–71. PMID: 10954255

18. Pandey M, Shukla VK, Singh S, Roy SK, Rao BR. Biliary lipid peroxidation products in gallbladder cancer: increased peroxidation or biliary stasis?. Eur J Cancer Prev 2000; 9: 417–22. PMID: 11201680

19. Shukla VK, Arya NC, Pitale A, Pandey M, Dixit VK, Reddy CD, Gautam A. Metallothionein expression in carcinoma of the gallbladder. Histopathology 1998; 33: 154–7. PMID: 9762548

20. Pandey M: Environmental pollutants in gallbladder cancer. J Surg Oncol 2006; 93(8):640–3 PMID: 16724354. 10.1002/jso.20531

21. Turgeon MK, Maithel SK. Cholangiocarcinoma: a site-specific update on the current state of surgical management and multi-modality therapy. Chin Clin Oncol. 2020 Feb;9(1):4. doi: 10.21037/cco.2019.08.09. Epub 2019 Sep 2. PMID: 31500433; PMCID: PMC7186525.

22. Zheng Q, Wu C, Ye H, Xu Z, Ji Y, Rao J, Lu L, Zhu Y, Cheng F. Analysis of the efficacy and prognostic factors of PD-1 inhibitors in advanced gallbladder cancer. Ann Transl Med. 2021 Oct;9(20):1568. doi: 10.21037/atm-21-4747. PMID: 34790774; PMCID: PMC8576663.

23. Roy N, Kshattry M, Mandal S, Jolly MK, Bhattacharyya DK, Barah P. An Integrative Systems Biology Approach Identifies Molecular Signatures Associated with Gallbladder Cancer Pathogenesis. J Clin Med. 2021 Aug 10;10(16):3520. doi: 10.3390/jcm10163520. PMID: 34441816; PMCID: PMC8397040.

24. Mishra SK, Kumari N, Krishnani N. Molecular pathogenesis of gallbladder cancer: An update. Mutat Res. 2019 Nov;816-818:111674. doi: 10.1016/j.mrfmmm.2019.111674. Epub 2019 Jul 6. PMID: 31330366.

25. Priya R, Pandey M, Shukla VK. Biomarkers in carcinoma of the gallbladder. Expert opinion on medical diagnosis 2008; 2:511–26.

26. Dixit R, Shukla VK, Pandey M. Molecular alterations in gallbladder cancer. World Journal of Pathology 2012, 1:7

27. Dixit R, Singh G, Pandey M, Basu S, Bhartiya SK, Singh KK, Shukla VK. Association of Methylenetetrahydrafolate Reductase Gene Polymorphism (MTHFR) in Patients with Gallbladder Cancer. J Gastrointest Cancer. 2016 Mar;47(1):55–60. doi: 10.1007/s12029-015-9794-0.

28. Srivastava V, Patel B, Kumar M, Shukla M, Pandey M. Cyclin D1, retinoblastoma and p 16 protein expression in carcinoma of the gallbladder. Asia Pacific J Cancer Prev Asian Pac J Cancer Prev. 2013; 14(5):2711–5

29. Dixit R, Pandey M, Tripathi SK, Dwivedi AN, Shukla VK. Comparative Analysis of Mutational Profile of Sonic hedgehog Gene in Gallbladder Cancer. Dig Dis Sci. 2017 Mar;62(3):708–714. doi: 10.1007/s10620-016-4438-1. Epub 2017 Jan 5. PMID: 28058596

30. .Maurya SK, Tewari M, Mishra RR, Shukla HS. Genetic aberrations in gallbladder cancer. Surg Oncol. 2012 Mar;21(1):37–43. doi: 10.1016/j.suronc.2010.09.003. Epub 2010 Sep 29. PMID: 20880699.

31. Dixit R, Pandey M, Tripathi SK, Dwivedi AND, Shukla VK. Genetic mutational analysis of β-catenin gene affecting GSK-3β phosphorylation plays a role in gallbladder carcinogenesis: Results from a case control study. Cancer Treatment and Research Communications, 2020; 23:100173 doi.org/10.1016/j.ctarc.2020.100173 PMID: 32344182

32. Liao Y, Wang J, Jaehnig EJ, Shi Z, Zhang B. WebGestalt 2019: gene set analysis toolkit with revamped UIs and APIs. Nucleic Acids Res. 2019 Jul 2;47(W1):W199-W205. doi: 10.1093/nar/gkz401. PMID: 31114916; PMCID: PMC6602449.

33. Xia J, Gill EE, Hancock RE. NetworkAnalyst for statistical, visual and network-based meta-analysis of gene expression data. Nat Protoc. 2015 Jun;10(6):823–44. doi: 10.1038/nprot.2015.052. Epub 2015 May 7. PMID: 25950236.

34. Szklarczyk D, Gable AL, Lyon D, Junge A, Wyder S, Huerta-Cepas J, Simonovic M, Doncheva NT, Morris JH, Bork P, Jensen LJ, Mering CV. STRING v11: protein-protein association networks with increased coverage, supporting functional discovery in genome-wide experimental datasets. Nucleic Acids Res. 2019 Jan 8;47(D1):D607–D613. doi: 10.1093/nar/gky1131. PMID: 30476243; PMCID: PMC6323986.

35. Franz M, Rodriguez H, Lopes C, Zuberi K, Montojo J, Bader GD, Morris Q. GeneMANIA update 2018. Nucleic Acids Res. 2018 Jul 2;46(W1):W60–W64. doi: 10.1093/nar/gky311. PMID: 29912392; PMCID: PMC6030815.

36. Korotkevich G, Sukhov V, Budin N, Shpak B, Artyomov MN, Sergushichev A. Fast gene set enrichment analysis. BioRxiv, 2021: 060012. https://www.biorxiv.org/content/10.1101/060012v3

37. Lo Surdo P, Calderone A, Cesareni G, Perfetto L. SIGNOR: A Database of Causal Relationships Between Biological Entities-A Short Guide to Searching and Browsing. Curr Protoc Bioinformatics. 2017 Jun 27;58:8.23.1-8.23.16. doi: 10.1002/cpbi.28. PMID: 28654729.

38. Hundal R, Shaffer EA. Gallbladder cancer: epidemiology and outcome. Clin Epidemiol. 2014 Mar 7;6:99–109. doi: 10.2147/CLEP.S37357. PMID: 24634588; PMCID: PMC3952897.

39. Dasari BVM, Ionescu MI, Pawlik TM, Hodson J, Sutcliffe RP, Roberts KJ, Muiesan P, Isaac J, Marudanayagam R, Mirza DF. Outcomes of surgical resection of gallbladder cancer in patients presenting with jaundice: A systematic review and meta-analysis. J Surg Oncol. 2018 Sep;118(3):477–485. doi: 10.1002/jso.25186. PMID: 30259519.

40. Haugsten EM, Wiedlocha A, Olsnes S, Wesche J. Roles of fibroblast growth factor receptors in carcinogenesis. Mol Cancer Res. 2010 Nov;8(11):1439–52. doi: 10.1158/1541-7786.MCR-10-0168. Epub 2010 Oct 13. PMID: 21047773.

41. Espinoza JA, Bizama C, García P, Ferreccio C, Javle M, Miquel JF, Koshiol J, Roa JC. The inflammatory inception of gallbladder cancer. Biochim Biophys Acta. 2016 Apr;1865(2):245–54. doi: 10.1016/j.bbcan.2016.03.004. Epub 2016 Mar 12. PMID: 26980625; PMCID: PMC6287912.

42. Chao TC, Wang CS, Jan YY, Chen HM, Chen MF. Carcinogenesis in the biliary system associated with APDJ. J Hepatobiliary Pancreat Surg. 1999;6(3):218–22. doi: 10.1007/s005340050110. PMID: 10526055.

43. Asai T, Loza E, Roig GV, Ajioka Y, Tsuchiya Y, Yamamoto M, Nakamura K. High frequency of TP53 but not K-ras gene mutations in Bolivian patients with gallbladder cancer. Asian Pac J Cancer Prev. 2014;15(13):5449–54. doi: 10.7314/apjcp.2014.15.13.5449. PMID: 25041017.

44. Hirata K, Kuwatani M, Suda G, Ishikawa M, Sugiura R, Kato S, Kawakubo K, Sakamoto N. A Novel Approach for the Genetic Analysis of Biliary Tract Cancer Specimens Obtained Through Endoscopic Ultrasound-Guided Fine Needle Aspiration Using Targeted Amplicon Sequencing. Clin Transl Gastroenterol. 2019 Mar;10(3):e00022. doi: 10.14309/ctg.0000000000000022. PMID: 30908307; PMCID: PMC6445609.

45. Liu P, Wang Y, Li X. Targeting the untargetable KRAS in cancer therapy. Acta Pharm Sin B. 2019 Sep;9(5):871–879. doi: 10.1016/j.apsb.2019.03.002. Epub 2019 Mar 6. PMID: 31649840; PMCID: PMC6804475.

46. Scanu T, Spaapen RM, Bakker JM, Pratap CB, Wu LE, Hofland I, Broeks A, Shukla VK, Kumar M, Janssen H, Song JY, Neefjes-Borst EA, te Riele H, Holden DW, Nath G, Neefjes J. Salmonella Manipulation of Host Signaling Pathways Provokes Cellular Transformation Associated with Gallbladder Carcinoma. Cell Host Microbe. 2015 Jun 10;17(6):763–74. doi: 10.1016/j.chom.2015.05.002. Epub 2015 May 28. PMID: 26028364.

47. Mohri D, Ijichi H, Miyabayashi K, Takahashi R, Kudo Y, Sasaki T, Asaoka Y, Tanaka Y, Ikenoue T, Tateishi K, Tada M, Isayama H, Koike K. A potent therapeutics for gallbladder cancer by combinatorial inhibition of the MAPK and mTOR signaling networks. J Gastroenterol. 2016 Jul;51(7):711–21. doi: 10.1007/s00535-015-1145-1. Epub 2015 Nov 27. PMID: 26614007.

48. Chen K, Zhu P, Chen W, Luo K, Shi XJ, Zhai W. Melatonin inhibits proliferation, migration, and invasion by inducing ROS-mediated apoptosis via suppression of the PI3K/Akt/mTOR signaling pathway in gallbladder cancer cells. Aging (Albany NY). 2021 Sep 27;13(18):22502–22515. doi: 10.18632/aging.203561. Epub 2021 Sep 27. PMID: 34580235; PMCID: PMC8507264.

49. Yang D, Chen T, Zhan M, Xu S, Yin X, Liu Q, Chen W, Zhang Y, Liu D, Yan J, Huang Q, Wang J. Modulation of mTOR and epigenetic pathways as therapeutics in gallbladder cancer. Mol Ther Oncolytics. 2020 Dec 3;20:59–70. doi: 10.1016/j.omto.2020.11.007. PMID: 33575471; PMCID: PMC7851494.

50. Wencong M, Jinghan W, Yong Y, Jianyang A, Bin L, Qingbao C, Chen L, Xiaoqing J. FOXK1 Promotes Proliferation and Metastasis of Gallbladder Cancer by Activating AKT/mTOR Signaling Pathway. Front Oncol. 2020 Apr 17;10:545. doi: 10.3389/fonc.2020.00545. PMID: 32363163; PMCID: PMC7180204.

51. Zong H, Yin B, Zhou H, Cai D, Ma B, Xiang Y. Inhibition of mTOR pathway attenuates migration and invasion of gallbladder cancer via EMT inhibition. Mol Biol Rep. 2014 Jul;41(7):4507–12. doi: 10.1007/s11033-014-3321-4. Epub 2014 Mar 13. PMID: 24623408.

52. Yang P, Javle M, Pang F, Zhao W, Abdel-Wahab R, Chen X, Meric-Bernstam F, Chen H, Borad MJ, Liu Y, Zou C, Mu S, Xing Y, Wang K, Peng C, Che X. Somatic genetic aberrations in gallbladder cancer: comparison between Chinese and US patients. Hepatobiliary Surg Nutr. 2019 Dec;8(6):604–614. doi: 10.21037/hbsn.2019.04.11. PMID: 31929987; PMCID: PMC6943012.

53. Amin M, Gao F, Terrero G, Picus J, Wang-Gillam A, Suresh R, Ma C, Tan B, Baggstrom M, Naughton MJ, Trull L, Belanger S, Fracasso PM, Lockhart AC. Phase I Study of Docetaxel and Temsirolimus in Refractory Solid Tumors. Am J Clin Oncol. 2021 Sep 1;44(9):443–448. doi: 10.1097/COC.0000000000000852. PMID: 34310349.

